# A passive protein environment offsets RNA folding energetics in cells

**DOI:** 10.64898/2026.09.07.749977

**Authors:** M. Todisco, F. Boskovic, A. Jain

## Abstract

RNA structure can be predicted from sequence in dilute solution, but these predictions often fail in cells. Many cellular RNAs keep their *in vitro* folds, whereas others are substantially less structured. We combined single-molecule FRET with *Xenopus* oocyte extract to measure RNA folding and duplex formation under cell-like conditions. Extract proteins passively suppressed base pairing: they slowed apparent strand association while leaving duplex dissociation largely unchanged, without sequence specificity or ATP consumption. A competitive binding model captured this activity as an effective folding penalty of ∼0.45 kcal/mol per nucleotide. Similar activity occurred in HeLa lysate. Applied to in-cell chemical probing data, this correction stratified reactivity across ∼42,000 stem-loops, whereas predictive models established in dilute solution classified nearly all as stably folded. Our results identify a passive protein-mediated mechanism as a key contributor to cellular RNA folding, shifting the balance between paired and unpaired states without changing the underlying base-pairing rules.

## INTRODUCTION

RNA folding shapes nearly every aspect of gene expression. The thermodynamics and kinetics of RNA base pairing are well described in dilute solution, where folding can be predicted from sequence using nearest-neighbor models (*1*, *2*). Applying these rules in cells, however, remains challenging. Methods to probe RNA structure in cells have revealed a mixed picture: many RNAs such as tRNAs (*3*) and viral IRES elements (*4*) adopt the structures expected *in vitro*, while broad regions of messenger RNAs are consistently less structured *in vivo* than expected from *in vitro* measurements and thermodynamic predictions (*5–7*). Likewise, aptamers and synthetic RNA devices that work *in vitro* often require sequence optimization before they fold in cells (*8*, *9*). Sequence-based prediction therefore holds for some cellular RNAs and fails for others, and what separates the two has not been established.

Reduced intracellular RNA structure is often attributed to energy-consuming remodeling by helicases, translating ribosomes, and other active processes; ATP depletion or translation inhibition, for example, increases mRNA structure (*5–7*). Yet active remodeling is only one feature of the intracellular environment and other components of the cellular milieu also affect RNA folding. Cellular metabolites and ions alter duplex stability (*10*), macromolecular crowding can favor compact states (*11–13*), and RNA-binding proteins may both stabilize (*14*) or disrupt (*15*) RNA structures without ATP hydrolysis. These factors act simultaneously and are difficult to tease apart in intact cells. Purified systems isolate individual contributions, but at the cost of losing the molecular complexity of the cytoplasm.

Cell-free extracts retain much of the molecular complexity of the cytoplasm while remaining accessible to biochemical manipulation. In particular, packed *Xenopus laevis* oocytes can be crushed by centrifugation without the use of lysis buffers, yielding a concentrated cytoplasmic layer with minimal dilution (Fig. 1A). *Xenopus* extracts have long been used to reconstitute complex cellular reactions in a cell-like environment (*16*). Here, rather than reconstructing a particular pathway, we used the extract itself as a tractable proxy for the cytoplasmic environment and asked how that environment alters RNA base pairing. The extract retains proteins, metabolites, and nucleic acids at near-physiological concentrations, yet remains amenable to dilution, fractionation by size or charge and enzymatic treatment. We combined this system with single-molecule microscopy, allowing us to follow intramolecular RNA folding and intermolecular duplex formation in real time within a cell-like environment.

**Fig. 1.**
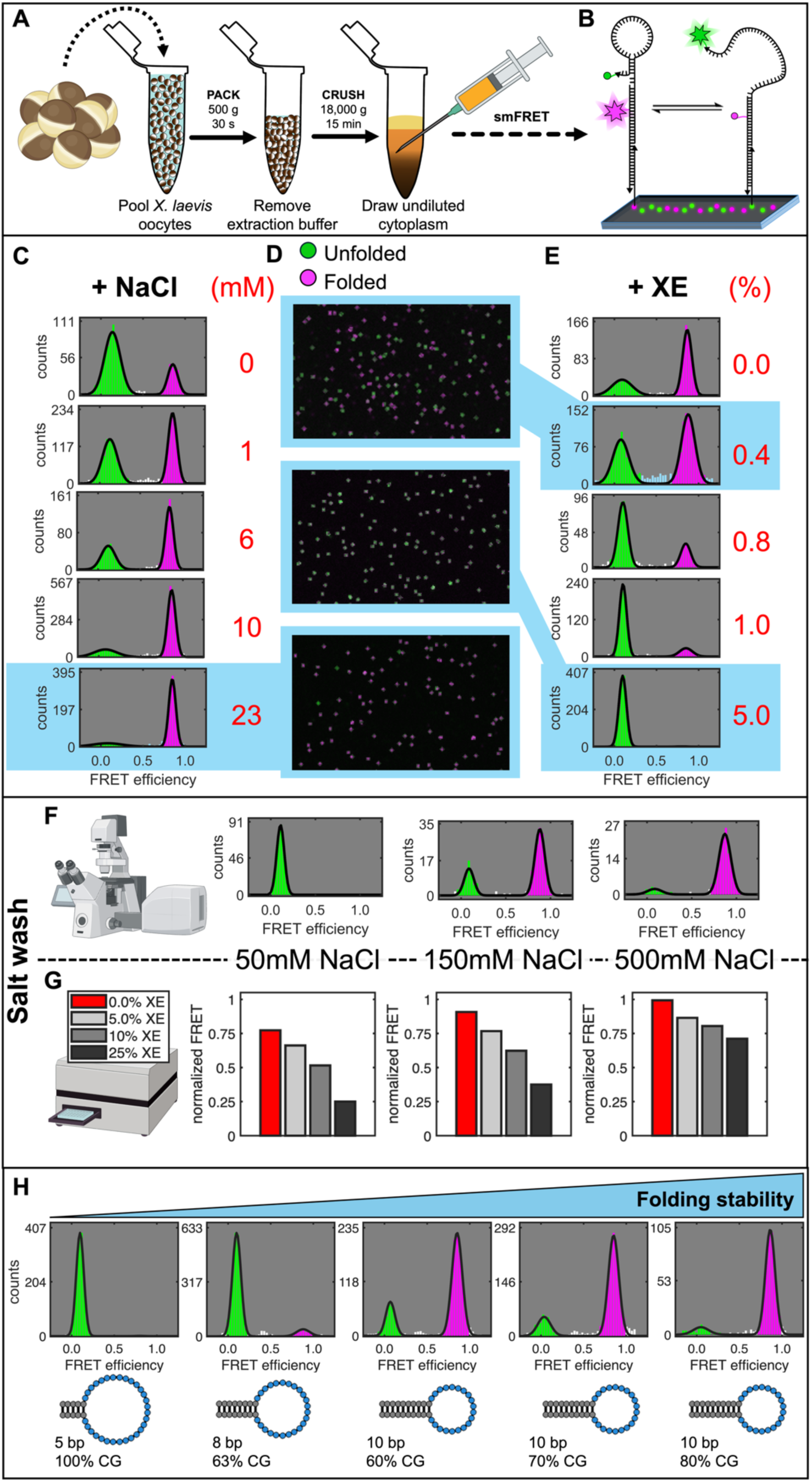
Xenopus extract reversibly destabilizes RNA hairpins. A) Workflow to extract the undiluted cytoplasm from *Xenopus laevis* oocytes. B) Single-molecule FRET assay used to monitor folding of surface-immobilized dual-labeled RNA hairpins. The observed color reports on the physical state of the construct. C) The construct progressively shifts from unfolded to folded state as salt is added to the buffered solution (3 mM Tris-HCl, pH 8.0). D) Representative micrographs show constructs as diffraction-limited spots acquired with smFRET. Green spots are interpreted as unfolded molecules, magenta spots are interpreted as folded molecules (see legend). E) In diluted buffer (7 mM Tris-HCl, pH 8.0) the construct mostly populates a folded high-FRET state. The addition of Xenopus Extract leads to greater population of unfolded molecules. F) Washing the microscope slide with low salt leaves the constructs unfolded once it has been exposed to Xenopus Extract. Washing the slide with high salt completely rescues folding. All samples were treated with 1.0% XE prior to salt wash. G) Ensemble measurements using a SpectraMax iD3 recapitulate the same observations from smFRET. Adding salt to our mixture reduces the effectiveness of XE, with unfolding becoming less impactful at very high salt. H) Hairpins predicted to have more stable folding energies also fold the best in XE. Thermodynamic stability still matters in the extract, but XE modulates folding.

Using this platform, we found that *Xenopus* extract shifts both RNA hairpins and bimolecular duplexes toward unpaired states. The effect arises primarily before helix formation: increasing extract concentration reduces the apparent rate of strand association by approximately fivefold, while leaving the dissociation rate of preformed duplexes nearly unchanged. Proteolysis abolishes the activity, whereas ATP depletion does not, and unrelated RNA, DNA, and polyphosphate titrate the activity away. These results support a passive mechanism in which cytoplasmic proteins bind exposed nucleic-acid segments and compete with helix formation.

Across a panel of simple stem-loops, folding depends jointly on conventional folding free energy and RNA length. A coarse-grained competition model represents this effect as an effective folding penalty of approximately 0.45 kcal/mol per nucleotide. The model predicts folding across a held-out hairpin series and is concordant with measurements in human lysate and structure-probing data from human cells, suggesting that protein competition raises the stability threshold required for sequence-encoded RNA structures to remain populated in cells. Because unrestrained base-pairing can drive widespread RNA association (*17*, *18*), we propose that this passive activity reflects a cellular strategy to keep the transcriptome soluble and accessible.

## RESULTS

### *Xenopus* oocyte extract suppresses RNA folding

To study the impact of the intracellular environment on RNA folding, we employed extract from *Xenopus laevis* oocytes (Fig. 1A). RNA folding was assessed using single molecule FRET. In brief, FRET donor (Cy3) and acceptor (Cy5) labeled constructs were sparsely immobilized on passivated microscope slides, and individual molecules were visualized as diffraction-limited spots (Fig. 1B). Constructs were designed such that the folded conformation resulted in high FRET between the two fluorophores. The system allowed facile exchange of the buffer, allowing us to tune and monitor RNA folding in real-time. Using a representative construct (RNA hairpin with 5 bp-long stem and 30 nt-long polyU loop), we found that adding Xenopus oocyte extract (hereafter XE) caused complete unfolding (Fig. S1, see Table S1 and Table S2 for list of sequences and constructs used). To improve the construct’s chemical stability, we then used a variant retaining the RNA stem and with a 30-nt polyT DNA loop (HP5). Surface immobilization did not alter its folding equilibrium (Fig. S2), and HP5 behaved consistently with standard thermodynamic predictions: it dynamically switched between open and closed states and was greatly stabilized by the addition of NaCl (Fig. 1C, D).

By titrating XE, we could continuously tune the environment from pure buffer (7 mM Tris, pH 8.0) to concentrated extract, and found that unfolding increased with XE concentration (Fig. 1D, E). Even at just 1.0% XE, the folded fraction of HP5 dropped from ∼75% to ∼15%, with a complete loss of measurable dynamics (Fig. S3). We observed comparable unfolding for a full-DNA construct with the same stem sequence (Fig. S1).

Importantly, this phenomenon was completely reversible, but only under the appropriate conditions. Upon washing with low salt, the construct remained unfolded. By contrast, upon washing with high salt (Fig. 1F), the original folded fraction and its dynamic behavior were fully restored, suggesting that salt is necessary to restore the folded state of the system and ruling out irreversible hydrolysis as an explanation for the observed unfolding.

The same qualitative behavior was reproduced performing ensemble measurements, showing reduced FRET with the addition of XE and a progressive rescue adding salt (Fig. 1G).

In buffer, hairpin folding is governed primarily by stem stability, set by stem length and GC content: longer, more GC-rich stems have more negative folding energies (ΔG) and higher folded fractions. We asked whether this rule still holds in extract. Although XE completely unfolded HP5, it grew progressively less effective against hairpins predicted to be more stable (Fig. 1H). Thermodynamic stability therefore still governs folding in the extract, with XE modulating the nearest-neighbor model rather than overriding it and motivating the development of a quantitative correction.

### Xenopus extract suppresses base-pairing by blocking strand association without sequence specificity

So far, we had examined intramolecular folding; we next asked whether the same destabilization extends to intermolecular hybridization (duplex formation). We set up a bimolecular RNA duplex assay in which a freely diffusing Cy5-labeled oligonucleotide hybridizes to a Cy3-labeled partner (Fig. 2A, B). In ensemble measurements, XE increased donor (Cy3) emission relative to the buffer control, indicating reduced duplex formation and mirroring the hairpin result (Fig. 2A).

**Fig. 2.**
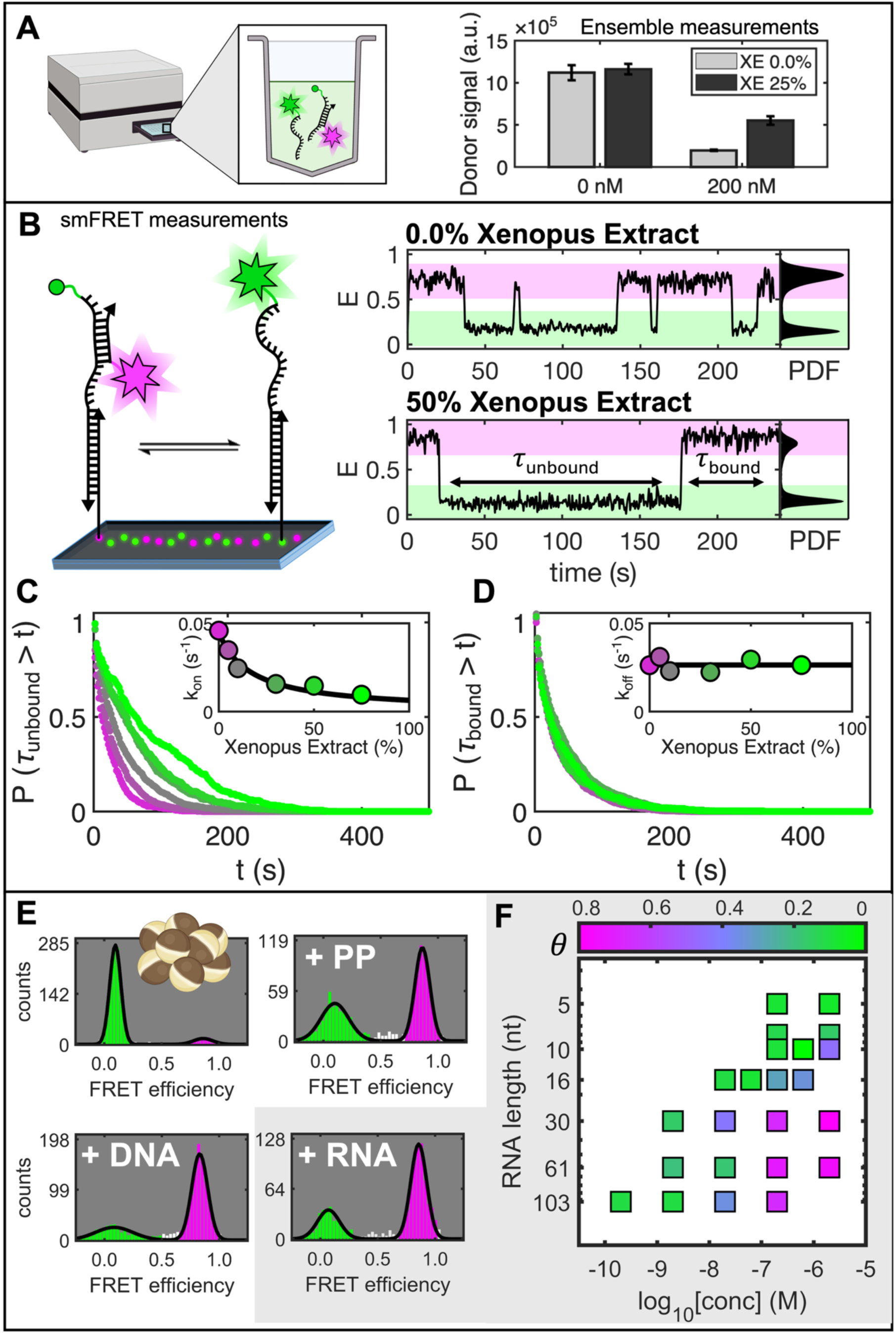
Xenopus extract suppresses RNA duplex formation by slowing strand association. A) Ensemble measurements of bimolecular RNA hybridization using a SpectraMax iD3 show that Xenopus Extract reduces the degree of duplex formation. Measurements performed with 50 nM of Donor (Cy3-labeled) RNA and 200 nM of Acceptor (Cy5-labeled) RNA in 50 mM NaCl, 10 mM Tris pH 8.0. B) Single-molecule assay for bimolecular RNA hybridization using smFRET with representative trajectories and state histograms in the absence and presence of Xenopus Extract. Histograms show probability density functions (PDF) for 0.0% and 50% XE, highlighting how XE shifts the population to single stranded. Measurements performed with 50 nM of Acceptor (Cy5-labeled) RNA in 50 mM NaCl, 10 mM Tris pH 8.0. C) Quantification of waiting-time distributions for duplex-formation events. Inset shows pseudo-first-order rates for *k_on_* decreasing in presence of XE. D) Waiting-time distributions for duplex-dissociation show little change in the first-order dissociation rate, indicating that the extract does not efficiently melt pre-formed duplexes. Inset shows first-order rates for *k_off_*. E) Competition experiments show that premixing the extract with unrelated polyanions, including ssDNA (14 nt, 200 nM), ssRNA (61 nt, 200 nM), and polyphosphate (PP, 140 monomers, 5 μM) restores hairpin HP5 folding as measured with smFRET. F) Results for rescue of HP5 folding with competitor RNAs having different lengths and concentrations. Symbols colors report the folded fraction of HP5 (*θ*, see colorbar) measured with smFRET. Folding of HP5 is rescued more effectively using longer and more concentrated competitors, consistent with nonspecific binding of the destabilizing activity to exposed nucleic-acid strands.

To further dissect this phenomenon, we turned to our smFRET assay by tethering the Cy3-labeled partner on the microscope slide (Fig. 2B). The same general behavior was reproduced there, with RNA hybridization reduced by XE (Fig. 2B, see histograms for XE 0.0% and XE 50%). However, smFRET allowed us to capture dynamic binding/unbinding of the two strands over time.

The destabilization mediated by Xenopus Extract could arise either because the extract actively melts base pairs that have already formed or because it prevents complementary strands from annealing in the first place. To distinguish between these mechanisms, we used time-resolved analysis. We reconstructed FRET trajectories as a function of XE, with probes transitioning back and forth from double helices (high FRET) to unpaired (low FRET) states (Fig. 2B). The waiting-time distributions for association (*τ*_unbound_, Fig. 2C) and dissociation (*τ*_bound_, Fig. 2D) events were consistent with a Poisson process characterized by a single rate constant.

Surprisingly, we found a highly asymmetric mechanism, with the dissociation lifetime of the duplex changing little across extract concentrations, indicating that XE does not efficiently accelerate melting of pre-formed duplexes. By contrast, the waiting time for association increased ∼5 times when going from pure buffer (50 mM NaCl, 10 mM Tris pH 8.0) to 75% XE, corresponding to a marked decrease in the apparent association rate. The simplest interpretation is that the extract acts primarily on single-stranded states and suppresses strand annealing before base pairs are formed. This mechanism is the opposite of what would be expected from helicase activity, known to speed up the dissociation rate of two strands (*19*).

If the same extract component that acts on single stranded RNA and prevents its hybridization is also responsible to suppress folding of the HP5 hairpin, then it should be possible to rescue its folding by titrating in single stranded RNA. Indeed, adding free unlabeled RNA restored HP5 folding (Fig. 2E) in a concentration-dependent manner (Fig. 2F). Moreover, single-stranded DNA and polyphosphate had a comparable effect (Fig. 2E), arguing that the interaction does not require RNA-specific chemistry.

This result suggested a potential lack of sequence specificity, as confirmed by a series of competition experiments with a panel of RNA sequences (Fig. 2F) with varying lengths (5–103 nt, see Table S1 for list of sequences). Surprisingly, this analysis revealed that longer competitor RNAs were much more effective than shorter ones (Fig. 2F), compatible with the existence of a defined binding footprint for the destabilizing factor.

Together with the salt reversibility of the activity, these observations support a largely electrostatic and sequence-independent interaction between RNA and a positively charged destabilizing factor in the extract.

### The folding-suppression activity resides in proteins and does not require ATP

The raw extract contains ∼ 45 g/l of proteins and ∼ 11 g/l of nucleic acid (see Materials and Methods), together with a rich ensemble of small molecules and ions, so we next asked what components of the extract were responsible for the suppression of base-pairing. Fractionation through a 3-kDa cutoff membrane separated the extract into a high-molecular-weight fraction and a light fraction enriched for small molecules and virtually void of proteins and nucleic acids. The light fraction was not competent in unfolding our HP5 construct (Fig. 3A). If anything, it modestly stabilized folding, consistent with an increase in ionic strength that favors RNA base pairing (Fig. 1C). Thus, the destabilizing activity is not explained by the small-molecule components of the extract alone.

**Fig. 3.**
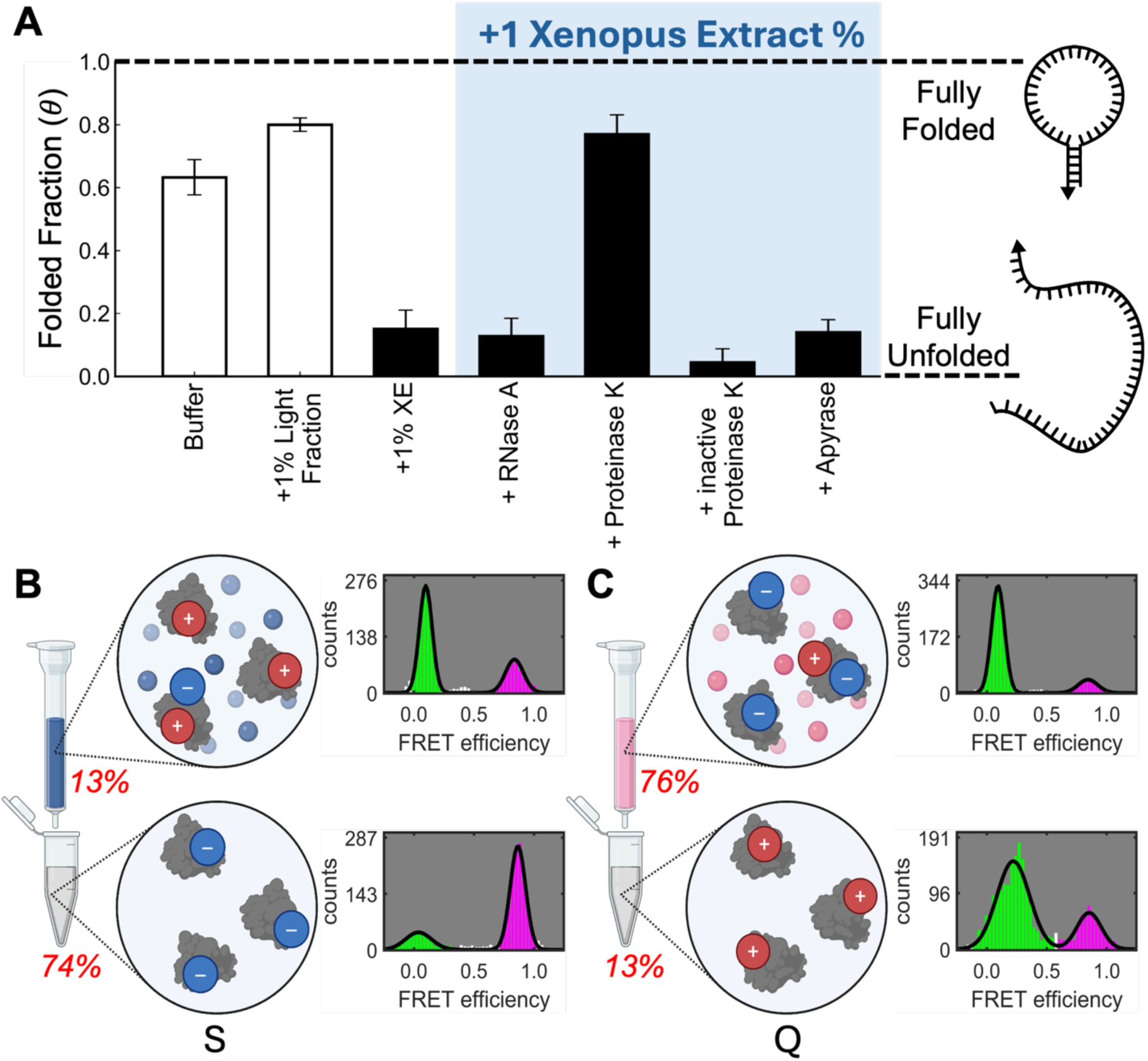
Folding suppression in Xenopus Extract is mediated by cationic proteins and ATP independent. A) Molecular weight fractionation of Xenopus Extract shows that losing the high-molecular-weight components causes the loss of unfolding activity (higher *θ*, folded fraction). Similarly, Proteinase K treatment abolishes hairpin unfolding, whereas RNase A or heat-treated Proteinase K do not. Apyrase also fails to restore folding despite the complete loss of ATP, indicating that the dominant anti-folding activity detected here is ATP independent and thus a passive process. B) Cation exchange fractionation of Xenopus Extract leads to an eluate (which retains ∼13% of the total proteins) which is competent in unfolding the HP5 hairpin. On the contrary, the flowthrough (containing ∼74% of the total proteins) loses its activity. C) Anion exchange fractionation of Xenopus Extract instead leads to an eluate (which retains ∼76% of the total proteins) and a flowthrough (containing ∼13% of the total proteins) that are both competent in unfolding the HP5 hairpin, suggesting that the presence of negative patches on the proteins does not allow to select for the unfolding proteins.

We next examined the role of high molecular weight components in attenuating RNA base pairing. To narrow down the potential candidates, we treated the raw extract with either RNase A or Proteinase K. Proteinase K treated extract lost its unfolding activity. In contrast, the folding of our construct was comparable in XE with or without treatment with RNase A or heat-inactivated Proteinase K. These results place proteins at the center of the RNA folding suppression mechanism.

We then asked whether ATP consumption was required, because ATP-dependent RNA remodeling has been invoked to explain reduced RNA structure *in vivo* (*5*). The raw Xenopus Extract contains ∼ 3 mM of ATP (see Fig. S4) which could fuel the unfolding activity. Apyrase treatment removed virtually all ATP from the extract (Fig. S4) but did not restore folding (Fig. 3A), showing that the dominant activity detected in our assay is ATP independent. The simplest model consistent with all these observations is that proteins in the extract bind exposed nucleic-acid strands and passively compete with base pairing.

The salt reversibility and competition experiments together pointed to an electrostatic interaction underlying this activity. Having established that proteins drive the unfolding, we asked whether they are cationic. Filtering XE through ion-exchange columns, we found that the vast majority of extract proteins are negatively charged, consistent with recent reports in Xenopus egg extract (*20*). The flowthrough of the cation (S) column (depleted of positively charged proteins) completely lost unfolding activity, whereas the eluate retained it (Fig. 3B), supporting positively charged proteins as the mediators. In contrast, both the eluate and flowthrough of the anion (Q) column were active (Fig. 3C), indicating that overall negative charge does not discriminate the unfolding proteins.

Together, these results indicate that accessible positive charges are required for the unfolding activity and provide a mechanistic rationale for its salt dependence (Fig. 1F, G). At low salt, the electrostatic interaction between these positive patches and the RNA backbone is strong, so the proteins remain bound even after the free extract is washed away, kinetically trapping the RNA in its unfolded state. At high salt, charge screening weakens the protein–RNA interaction and releases the RNA, allowing it to refold.

### A competitive binding model yields a predictive per-nucleotide correction

Our study on hairpin HP5 revealed a dramatic unfolding of RNA in Xenopus Extract. This result is puzzling because the intracellular environment must maintain the many structured RNAs essential for life. Some RNAs must therefore meet specific conditions that permit folding despite this activity.

To identify these conditions and convert them into a quantitative framework, we measured a panel of 10 hairpins across a range of Xenopus Extract concentrations. The constructs spanned total lengths of 15-40 nucleotides, with 4-10 bp stems and 5-30 nt loops. Their predicted folding energy (160 mM NaCl, 20°C) ranged from -4.5 kcal/mol to -23.9 kcal/mol (Fig. S5 and supplementary materials give the salt calibration for the extract). If the extract behaved as a simple buffer, all of them would be fully folded in pure XE.

Across the panel, we found that not all hairpins were unfolded by Xenopus Extract. Instead, the constructs displayed a range of behaviors, from monotonic unfolding to non-monotonic profiles (Fig. 4A-D). Because extract simultaneously contributes protein-mediated destabilization and ionic-strength–dependent stabilization, this behavior reflects the complex interplay between the two. We do not attempt to model this intermediate regime, as RNA thermodynamics at the low ionic strengths sampled during the titration is poorly parameterized; we therefore restrict our considerations and quantitative analysis to the folded fraction extrapolated to pure extract, the relevant endpoint representing the conditions of the intracellular environment.

**Fig. 4.**
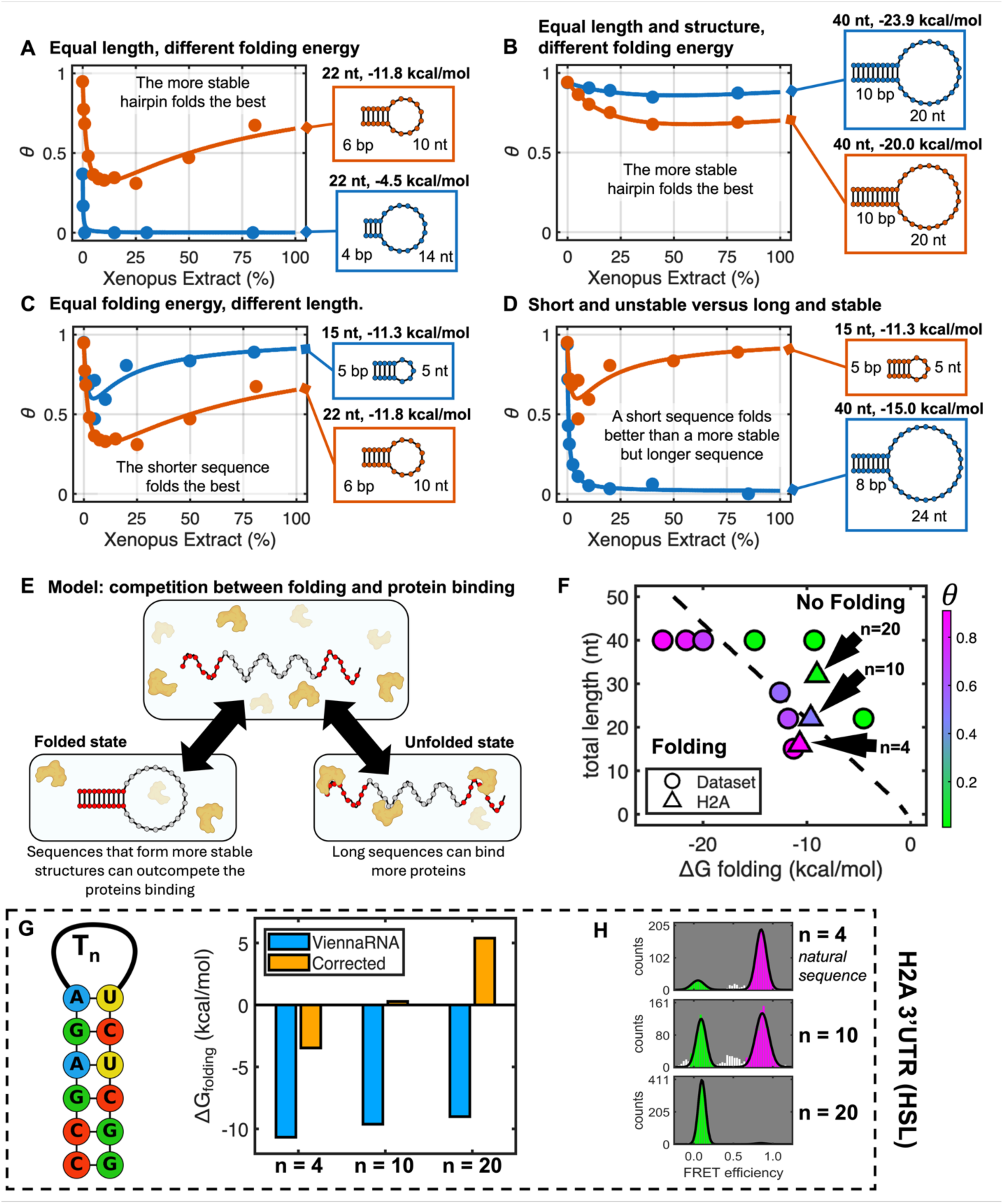
Extract-mediated RNA unfolding is a function of folding energy and RNA length. A, B, C, D) Comparison of folded fractions (*θ*) measured with smFRET at various Xenopus Extract concentration for a panel of hairpins having various lengths and thermodynamic stabilities. Sketches depict the measured hairpins and state their total length (nt), stem length (bp), loop length (nt) and predicted folding energy (160 mM NaCl, 20°C). A) Comparison between two constructs having same total length (22 nt) but different folding stability. Although both are predicted to fold in the intracellular environment, only the more stable one folds in XE. B) Comparison between two constructs having same total length (40 nt) and structure (same stem and loop lengths) but different CG-content. Both constructs fold as expected in the intracellular environment, but the more stable one folds the best. C) Comparison between two constructs having same folding energy (∼ -11.5 kcal/mol) but different lengths and structure. Both constructs fold as expected in the intracellular environment, but the shortest one folds the best. D) Comparison between a relatively unstable but short construct and a more stable but long construct. Although both are predicted to fold in the intracellular environment, only the short construct folds in XE. E) Sketch for proposed model of Xenopus Extract mediated destabilization of RNA folding. The unfolded RNA strand binds to proteins that prevent folding. Longer sequences will bind more proteins and thus being more vulnerable. If the folded state is stable enough, the protein binding can be outcompeted to restore structure. F) Scatter plot for the folded fraction in pure XE of the hairpins studied here (circles) as a function of their computed folding energy (160 mM NaCl, 20°C) and total length. Dashed line shows the boundary separating folded and unfolded hairpins according to our model. Triangles show validation data for H2A 3’UTR, not used to establish the model. G) Comparison of standard folding predictions for human H2A 3’UTR sequence (HSL) and its variants with the corrected in-cell energies inferred from this work. While ViennaRNA always predicts stable folding, the corrected model suggests that the unnatural variants should not fold fully in the intracellular environment. H) Folded fraction of HSL in 80% XE for natural sequence and its variants having longer loops. The folded fraction closely tracks predictions from our model (see Fig. 4G, F).

Focusing on this endpoint, we first asked whether very stable hairpins might escape unfolding. Comparing two constructs of equal length (22 nt), we found that the more stable stem-loop folded better in Xenopus Extract (Fig. 4A).

Because these two sequences formed structures that differed slightly in stem and loop length, we next removed those variables by comparing two 40-nt constructs, each with a 10-bp stem and a 20-nt loop, differing only in stem GC-content (Fig. 4B). The construct with higher GC-content (and thus more stable folding) again folded better. Greater thermodynamic stability therefore attenuates the unfolding activity of the extract.

Our competition experiments (Fig. 2F) had hinted that longer RNAs are more vulnerable to the activity. To isolate length, we studied two sequences of comparable folding energy but different length (Fig. 4C). Strikingly, the longer sequence was more heavily destabilized in Xenopus extract, despite being predicted to form a slightly more stable hairpin in pure buffer.

Together, these observations point to a system governed by two variables: RNA folding energy and construct length, in which only very stable or very short stem-loops resist the unfolding activity. This raises a counterintuitive prediction: a short hairpin could fold better than a longer but more stable one. Testing this directly, we compared a very short hairpin (15 nt) with a longer, more stable one (40 nt); only the short construct folded in the extract (Fig. 4D).

Combining these results with the outcome of our biochemical manipulation (Fig. 3A), we propose a simple model where long single-stranded RNA sequences bind many proteins that block folding. This effect can be outcompeted if the RNA folding energy is large enough to offset the protein binding energy (Fig. 4E).

Plotting the folded fraction against the model’s two parameters (Fig. 4F), we found that hairpins folding in Xenopus Extract cluster in the region combining short length and stable stems. We formalized this as a direct competition, in which the open hairpin can either close into a stem-loop or bind the extract-derived proteins (see supplementary materials for derivation). This model is controlled by a single free parameter describing the *per-nucleotide* binding energy of the protein 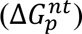.

With a protein-binding energy 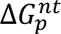 of ∼ -0.45 kcal/mol per nucleotide, the model splits the dataset into two regions (Fig. 4F, dashed line): one where folding is permitted (stable stems, short sequences) and one where it is not (weak stems, long sequences), recapitulating the measured behavior.

Crucially, the model also generalizes to sequences outside our training set. Beyond reporting the folded fraction of a given hairpin, it corrects the predicted folding energy of arbitrary sequences. The protein-binding energy 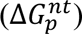 it defines acts as an effective energetic penalty on folding (see supplementary materials). In other words, the intracellular environment shifts the apparent folding free energy by a net ∼ 0.45 kcal/mol per nucleotide toward less favorable values. This offset can be applied as a correction to standard tools such as ViennaRNA, NUPACK, and mfold, which consistently overestimate folding under cell-like conditions (Fig. 4G).

To test this, we turned to the histone mRNA 3′UTR stem-loop (HSL), a structured element that binds SLBP and is required for downstream processing of the transcript. This 16-nt element folds into a short 6-bp stem capped by a 4-nt loop. We applied our model to the human H2A HSL and to variants with progressively longer loops. ViennaRNA predicted all of them to remain fully folded. Our correction, by contrast, predicted full folding only for the native HSL (corrected energy −3.5 kcal/mol); the 10-nt-loop variant (22 nt, +0.3 kcal/mol) should be partially folded, and the 20-nt-loop variant (32 nt, +5.4 kcal/mol) fully unfolded (Fig. 4G).

This prediction was validated using our smFRET assay (Fig. 4H, 4F), confirming that only the native sequence was fully folded as required by its biological function, with longer sequences progressively unfolding, in line with the corrected energies.

Thus, the main contribution of the model is not merely descriptive; it provides a practical way to adjust RNA-folding predictions for the intracellular environment.

### The competitive model improves prediction of human-cell RNA structural data

Next, we asked whether the observed phenomenon was due to a protein component specific to *X. laevis* and/or its oocytes. A commercial HeLa lysate unfolded HP5 in the same qualitative manner as XE. After normalizing by total protein concentration, the potency of HeLa lysate was only slightly lower than XE, supporting the idea of a shared common mechanism in the intracellular milieu rather than a Xenopus-specific factor (Fig. 5A).

**Fig. 5.**
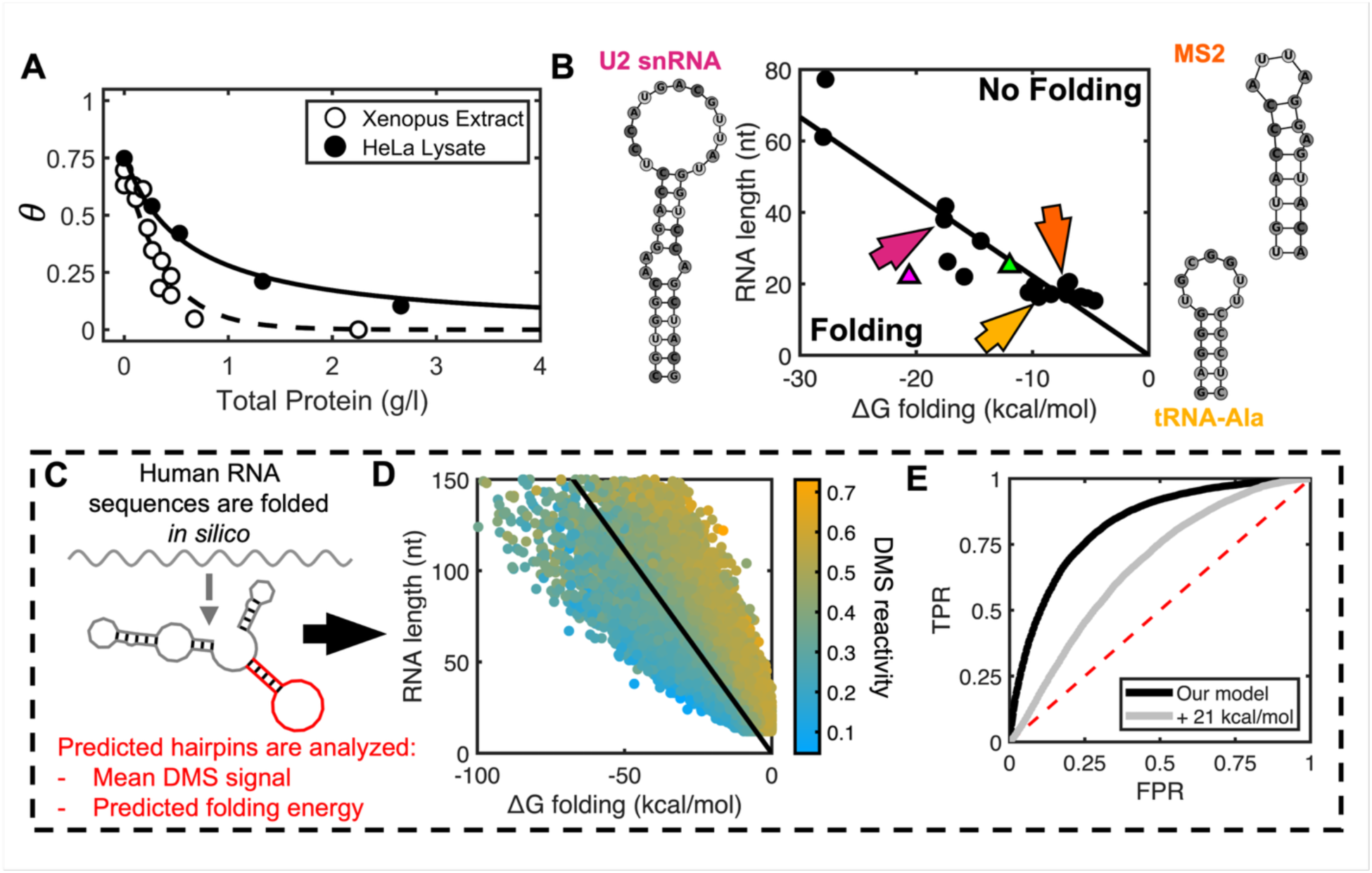
Model validation for human-cell RNAs. A) HeLa lysate causes HP5 hairpin to unfold. After normalizing for the total protein content (∼27 g/l in HeLa lysate vs ∼ 45 g/l in XE), human extract shows only a slightly lower potency in destabilizing RNA hairpins, supporting a shared protein-dependent mechanism. B) Curated dataset of RNA structures known to fold in human cells is shown against our model. Solid black dots represent sequences whose minimum free energy structure matches the one from RNAcentral and can be thus considered thermodynamically determined. RNA structures completely avoid the top-right region of the graph as predicted by our model. Triangles represent Spinach SL3 (green) and the highly fluorescent Superfolding Spinach SL3 (magenta) from Strack *et al*. Solid line shows our model parametrized according to our results with Xenopus Extract. C) Schematics for the retrieval and computational analysis of stem-loop domains from human mRNA sequences. D) Distribution of DMS reactivities in the whole dataset (mRNA UTRs, ∼42000 hairpins) matches expectations from our model (solid line). DMS reactivities have been smoothed for visualization purposes. E) AUC ROC for DMS reactivity vs folded state computed with our model (black) or using a flat correction to ViennaRNA energies (gray) of + 21 kcal/mol to obtain matching folded fractions. The folded/unfolded assignments predicted by our model are more concordant with the measured DMS reactivities than those from ViennaRNA. Dashed red line shows random expectation.

By simple thermodynamics, any RNA domain with a predicted folding energy of just a few kcal/mol should be fully folded inside the cell. Instead, the protein activity we characterized imposes a boundary on which sequences can fold in the intracellular environment, resolving our initial question.

Because the penalty scales with length, an RNA structure persists in cells only when its folding stability outpaces this per-nucleotide cost: highly stable natural elements clear the threshold despite their size and fold much as they do *in vitro*, whereas short, only moderately stable helices or having long loops would fall on the unfolded side — explaining why designed elements so often require extensive tuning to function in cells (*8*, *9*). This conclusion prompted us to check if a panel of sequences known to fold inside the human cell follow this rule. To answer this question, we manually curated a dataset of 18 RNA stem-loops belonging to different families including tRNA, snRNA, pre-miRNA and scaRNA, and computed the folding energy (using ViennaRNA) of their structure as reported by RNAcentral (*21*) (see Table S3). For this study we focused on those sequences whose structure was in agreement with the minimum free energy structure as determined with ViennaRNA – and can thus be considered thermodynamically determined. All 18 sequences avoided the top-right region (Fig. 5B, black dots), falling near the boundary and slightly toward the folding-permissive side. This small offset is in the direction expected from the (i) higher temperature of human cells compared to our experimental conditions and (ii) the modestly lower unfolding potency of human cells (Fig. 5A), consistent with the Xenopus-derived boundary being a conservative estimate.

On top of these structured sequences, we also verified the stem-loop 3 (SL3) that was re-engineered by Strack *et al.* (*8*) – among other changes – to transform the Spinach aptamer into its superfolder variant (Spinach2) that works more efficiently in cells. Consistent with our model, the parent Spinach SL3 sits close to our boundary, whereas the stabilized Spinach2 SL3 lies well within the folded region (Fig. 5B, triangles).

We then asked whether the same framework could explain transcriptome-wide RNA-structure measurements in human cells. Using published DMS-TRAM-seq data from U-2OS cells (*7*), we extracted ∼42,000 predicted stem-loop domains (*2*) from the UTRs of endogenous RNAs and computed the average DMS reactivity of their stems together with their folding energy (Fig. 5C). Importantly, almost all hairpins (41735 out of 41847) retrieved this way were predicted to be stably folded by ViennaRNA, meaning that DMS reactivity could not be used as a predictor for their physical state within the conventional thermodynamic framework.

In the “folding energy vs length” space, we found that the folded domains’ reactivities tracked our model closely, with stem-loops having high reactivity (likely unfolded) in the top right region of our graph and stem-loops having low reactivity (likely folded) in the bottom left region of our graph (Fig. 5D).

To quantify whether DMS could effectively discriminate between the folded/unfolded state according to our model, we split the dataset into sequences predicted to be folded (*θ* > 0.5; 10620 entries) or unfolded (*θ* < 0.5; 31227 entries) and computed the AUC for the ROC (Receiver Operating Characteristic), finding an excellent score of 0.83 (Fig. 5E).

ViennaRNA classifies almost all hairpins as folded, indicating a systematic overestimation of RNA folding stability in the cell. This raises a concern: our model might outperform ViennaRNA simply because it destabilizes RNA broadly, rather than because it captures the specific destabilization mechanism. To distinguish these possibilities, we tested our model against a null model: ViennaRNA destabilized by a flat energy correction. We increased this correction until it yielded the same fraction of folded domains as our model (Fig. 5E). At this matched folded fraction, our model was significantly better, reaching AUC ∼0.83 versus ∼0.67 for the flat correction (+21 kcal/mol; ΔAUC ∼0.16, 95% CI [0.152, 0.164], p < 2×10⁻⁴, paired bootstrap over N = 41,847). A flat correction unfolds domains according to stability alone, but these are not the domains that DMS reports as unfolded, so it cannot match our model’s agreement with the data. The improvement therefore arises from the specific, length-dependent form of our correction, not from destabilization alone.

These results argue that the passive destabilization measured biochemically is not an idiosyncrasy of Xenopus extract but generalizes to human (HeLa lysate, curated folded RNA dataset, U2OS UTR DMS data), potentially reflecting a general feature of intracellular RNA folding.

## DISCUSSION

By combining sm-tFRET with computational analysis, we dissected the behavior of RNA base-pairing inside *Xenopus* extract. Studying a series of RNA hairpins, we found that base-pairing is strongly destabilized in the intracellular environment, in qualitative agreement with earlier reports (*5–7*). Fractionation and enzymatic treatments revealed that proteins are responsible for this effect. Most notably, the destabilization is due to an electrostatically mediated interaction that is not specific to RNA, as it also affects other negatively charged polymers such as DNA and polyphosphates. Surprisingly, this phenomenon does not depend on ATP consumption, suggesting a mechanism entirely distinct from and parallel to the ATP-dependent remodeling previously described in yeast by Rouskin and colleagues (*5*), at the same time excluding helicase activity as a potential candidate.

The discrepancy between *in vitro* folding energies and in-cell structure measurements has often been treated qualitatively, but our data show that it can be formalized as an energetic correction. For short and moderately stable stem-loops, standard folding tools are systematically too optimistic about the persistence of RNA structure in cells. Incorporating a passive protein-mediated penalty of ∼ 0.45 kcal/mol per nucleotide, we substantially improved the agreement with both our single-molecule measurements and transcriptome-wide human data. This provides a practical route toward more realistic prediction of intracellular RNA folding and offers a quantitative explanation for why many designed RNAs work well *in vitro* yet fail in cells unless heavily optimized for stability (*8*).

Conceptually, this places our mechanism in opposition to molecular crowding: excluded-volume effects from inert macromolecules have been reported to stabilize intramolecular interactions and folding (*11*, *12*, *22*, *13*), whereas the protein environment that we characterized opposes base pairing. Importantly, since the mechanism presented here relies on protein binding, our correction has to be applied both to pairing and non-pairing nucleotides, making it fundamentally different from other corrections to the nearest neighbor parameters that have been independently proposed by other groups to account for *in vivo* conditions (*10*, *23*, *24*).

These findings raise the question of why cells require such a broad passive strategy to suppress RNA base-pairing. One possibility is that keeping RNA unfolded facilitates regulation and ensures that sequences relevant for biological pathways remain exposed and accessible. This may be critical because RNA base-pairing is intrinsically strong: it can drive widespread aggregation of purified mRNAs (*17*, *18*) and promote intracellular aggregates (*25*) such as stress granules (*17*, *26*), TIS granules (*27*), and RNA-repeat disorder aggregates (*28*, *29*).

We can therefore speculate that this destabilization mechanism has evolved to prevent RNA aggregation and maintain the transcriptome in a functional state (*18*). Such process would echo the activity of known and much more specific proteins such as CspA, which disrupts RNA duplexes during cold shock (*30*), and Staufen 1, which resolves base-pairing between mRNAs and lncRNA Alu elements (*31*).

Our approach carries several limitations that also mark clear directions for future work. First, although undiluted oocyte extract preserves the macromolecular composition of the cytoplasm at near-physiological concentrations, it is not an intact cell: compartmentalization, active transport, and ongoing transcription and translation are absent. The quantitative agreement with human in-cell DMS data (Fig. 5) nonetheless argues that the dominant activity we measure also operates in living cells. Second, all of our thermodynamic analysis relies on the folded fraction extrapolated to pure extract, as the intermediate-concentration regime convolves protein-mediated destabilization with ionic-strength effects that we deliberately do not model. Third, the ∼0.45 kcal/mol per nucleotide penalty is an effective parameter: because the protein footprint and the per-site binding energy cannot be determined independently from our data (see supplementary materials), the correction is best regarded as a phenomenological rule rather than a microscopic binding constant. Resolving this would require identifying the dominant destabilizing proteins, together with understanding the extent to which this activity is distributed across many abundant RNA-binding proteins versus concentrated in a smaller subset of specialized proteins. Finally, our calibration and validation rest on simple stem-loops at a single reference temperature; extending the framework to multi-helix architectures, tertiary contacts, and the higher temperatures of mammalian cells (where we already detect a small offset, Fig. 5A, B) will be needed to turn this correction into a general predictive tool.

In summary, our work uncovers a protein-mediated, sequence-independent suppression of RNA folding rooted in the physical chemistry of RNA. This mechanism helps reconcile *in vivo* and *in vitro* measurements and offers a quantitative route to more realistic folding predictions. More broadly, understanding how cells control the physical state of RNA may illuminate transcriptome organization and point to therapeutic opportunities in diseases of RNA misfolding and aggregation.

## MATERIALS AND METHODS

### Oligonucleotides synthesis and purification

All DNA oligonucleotides were obtained from IDT with provided HPLC purification. RNA oligonucleotides were synthesized in-house on an H-6 K&A solid-phase nucleic acid synthesizer, following the manufacturer recommended protocol. The protected oligonucleotides were cleaved from the solid support using a 1:1 v/v mixture of ammonium hydroxide (30% NH_3_ in water) and aqueous methylamine. The cleaved material was heated up to 65°C for 15 min to deprotect the nucleobases. Ammonia and methylamine were removed by evaporation in a vacuum centrifuge, and the leftover water was removed through lyophilization. To remove the 2′-OTBDMS-protecting groups, the samples were dissolved in 100 μL of DMSO and 125 μL of TEA.3HF and then heated to 65 °C for 2.5 h. The oligonucleotides were precipitated adding a mixture of 0.1 V of ammonium acetate and 5 V of isopropanol for 20 min at −80 °C. The oligonucleotides pelleted this way were then washed once with 80% ethanol and dissolved in 100 μL of formamide. The collected material was further purified by denaturing 20% PAGE. The full-length products were cut out, crushed, and soaked for 16 h in 5 mM sodium acetate pH 5.5 and 2 mM EDTA pH 8. The oligonucleotides were then desalted using C18 cartridges from Sep-Pak (Waters) and lyophilized. Reagents and consumables for oligonucleotides synthesis and purification were acquired from ChemGenes and Glen Research. Reagents for cleavage and deprotection were acquired from MilliporeSigma.

### Preparation of Xenopus Extract

The defolliculated *Xenopus laevis* oocytes were acquired by Xenopus 1 Corp and immediately processed upon receival. The oocytes were kept on ice during the whole procedure and extensively washed with 4 L of extraction buffer (*32*), consisting of 10 mM HEPES pH 7.5, 10 mM sodium acetate, 1 mM magnesium acetate and 2 mM DTT. This process gets rid of debris and enriched the sample in large and healthy oocytes. Leftover small or anomalous oocytes were manually removed using a transfer pipette. The oocytes prepared this way were transferred to 13 mL centrifuge tubes for the packing and crushing steps. All centrifugations were performed with equipment set to 4°C.

The packing step was performed at 500g for 30 seconds in an Eppendorf Centrifuge 5810R. The supernatant was removed and the step repeated once again to remove as much buffer as possible and avoid diluting the final extract. The tubes were then transferred to a Beckman Coulter Avanti JXN-26 high-speed centrifuge using the fixed-angle rotor model JA-25.50. The samples were spun at 18,000g for 15 minutes. This process broke open the oocytes and stratified their content in lipid-rich layer (top), cytoplasm layer (middle) and pigment layer (bottom). The middle layer was carefully drawn by piercing the wall of the tube with a syringe and transferred to 1.5 mL tubes. These tubes underwent a final clean-up step by briefly spinning them on a fixed-speed benchtop centrifuge before once again drawing the clear middle layer by piercing the wall with a syringe, and aliquoted for long-term storage.

The aliquots of extract prepared this way were flash-frozen in liquid nitrogen and stored in -80°C freezer. Whenever used, they were thawed and kept in ice.

### Fractionation of Xenopus Extract

To obtain the light fraction of the Xenopus Extract, we spun the raw extract in Amicon Ultra 3K filters (MilliporeSigma) for 15 min at 15,000 x g and collected the flowthrough. The light fraction had no detectable proteins and ∼ 1 g/l of residual nucleic acid.

Ion exchange for protein fractionation was performed using Mini Pierce Strong Ion Exchange Columns (4 mg capacity, Thermo Fisher Scientific). The columns were conditioned with 25 mM Tris-HCl pH 8.0, and the Xenopus Extract sample was loaded after dilution (1:6) in 10 mM Tris-HCl pH 8.0. After loading, the column was washed with 25 mM Tris-HCl pH 8.0 before final elution in 25 mM Tris-HCl pH 8.0, 2 M NaCl.

To clean up after the ion exchange, we used Micro Bio-Spin™ P-30 Gel Columns (BioRad) following instructions from the manufacturer. The columns were pre-equilibrated with Tris-HCl 10mM pH 8.0, 150mM NaCl.

### Microscopy samples preparation

The physical state of nucleic acids was determined using sm-tFRET. The samples were loaded in flow cells assembled between a standard microscope slide (Fisher Scientific plain premium microscope slides) and a coverslip (VWR micro cover glass) held together by double-sided tape. To load the samples, two holes were drilled on the slide using a Dremel 200 rotary tool with diamond-coated drill bit, and the two glasses were sealed together using dual-component 5 minutes epoxy (Devcon). The slides and coverslips were both coated with PEG to reduce non-specific interactions, and the coverslips were functionalized with biotinylated PEG to allow our construct to bind.

Microscope slides were treated as follows: (i) slides and coverslips were places in Coplin jars and sonicated with DI water for 10 minutes, followed by (ii) sonication with Methanol for 10 minutes and (iii) sonication with 2 M Potassium Hydroxide for 20 minutes. After washing with Methanol, (iv) the glass was allowed to react with with 1 % *N*-[3-(Trimethoxysilyl)propyl]ethylenediamine (MilliporeSigma 104884) in Methanol for 10 minutes in the dark, followed by a brief sonication and 10 more minutes on reaction in the dark. After the reaction was completed, the slides were washed with Methanol and fully dried in an oven set at 50°C. When completely dried, (v) each microscope slide was allowed to react with 16 mg of mPEG-SVA-5000 (Laysan Bio) in 70 μL of 100 mM Sodium Bicarbonate, and each coverslip was allowed to react with 16 mg of mPEG-SVA-5000 + 0.3 mg of Biotin-PEG-SVA-5000 (Laysan Bio) in 70 μL of 100 mM Sodium Bicarbonate for 3 h at room temperature. (vi) After reaction completion, the slides and coverslips were thoroughly washed with milliQ water and dried under air stream before being vacuum sealed and stored at -20°C.

To physically tether the nucleic acids to the microscope slides, we employed a multi-stranded design (see Table S1 and Table S2). The sequences were pre-mixed at a final concentration of 5 μM in T50 buffer (50 mM NaCl, 10 mM Tris pH 8.0), heated up to 75°C and then cooled down over 15 minutes to anneal. To allow the coverslips to bind the constructs, the flow chambers were primed with T50 buffer and then allowed to react with 0.2 g/L of NeutrAvidin (Invitrogen) in T50 for 15 minutes before being washed with T50. The constructs were loaded and allowed to bind to the biotin/neutravidin complexes for 10 minutes before flowing with the imaging buffer.

### sm-tFRET measurements

All measurements were performed in imaging buffer containing Glucose Oxidase from *Aspergillus niger* (100 U/mL), Catalase from bovine liver (1500 U/mL), Trolox (3 mM) and D-(+)-Glucose (0.8% w/v). All enzymes and chemicals were acquired from MilliporeSigma.

Measurements for the intermolecular hybridization of RNA duplexes was performed by supplementing 10 mM EDTA, 10 mM EGTA and 40 U of RNase inhibitor.

Single molecule measurements were performed with a Nikon Eclipse Ti inverted microscope equipped with an iLas ring-TIRF controller (Gataca Systems). The samples were imaged using a 532 nm Cobolt laser (Hübner photonics) through a 100x oil-immersion TIRF objective. Using the Gemini W-view (Hamamtsu), the light collected from the objective was split in its short-and long-wavelength components with a 600 nm dichroic mirror before being imaged side by side onto two separate parts of the iXon Ultra EMCD Camera sensor (Andor) allowing for the imaging of the two channels simultaneously. The two channels were spatially registered using a 4-th degree polynomial transformation to align the signals coming from fluorescent polysterene tracers (FluoSpheres red 0.1 μm, Thermo Fisher Scientific). Data was collected using μManager (*33*) in lossless TIFF format and analyzed with custom MATLAB code to identify fluorescent particles and record their signal intensity. To extract transition rates and dwell times, FRET trajectories for each condition were globally fitted using the Viterbi algorithm (*34*) and assuming a shared underlying Hidden Markov Process as described in literature (*35*).

### Enzymatic treatment

To evaluate the role of different intracellular components on the unfolding capability of the Xenopus Extract, we performed a series of enzymatic treatments. For the Proteinase K treatment, 10 μL of Proteinase K were added to 100 μL of XE and left reacting at room temperature for 2 h (final concentration 1.8 g/L). As a control, the Proteinase K was pre-heated at 95°C for 15 minutes to inactivate it before adding it to the XE.

For RNase A treatment, a 1:20 dilution was prepared, and 1 μL of this solution were pre-diluted in 10 μL milliQ water and then added to 50 μL of XE and allowed to react for 2 h at room temperature (final concentration 8 mg/L). As a control, 1 μL of the diluted RNase A was pre-mixed with 10 μL of RNase inhibitor before being added to XE. After two hours at room temperature, 10 μL of inhibitor were added to the first vial before performing our measurements (final concentration of inhibitor 7 U/μL).

For the Apyrase treatment, 2 μL of the enzyme were added to 50 μL of XE and allowed to react for 2 h (final concentration 20 U/L).

Proteinase K (25530049), RNase A (EN0531) and RNase inhibitor (EO0381) were acquired from Thermo Fisher Scientific. Apyrase was bought from New England Biolabs (M0398S).

### ATP quantification and Apyrase activity assessment

To quantify the amount of ATP in the extract, we used the ATP-Assay Kit-Luminesce from Dojindo (product A550). The assay reports on Luciferase activities and relies on the sample’s ATP to fuel the reaction and convert D-Luciferin to Oxyluciferin, emitting visible light as a byproduct. A calibration curve was measured in triplicate as recommended by the producer, and the light emitted by XE treated with either added Apyrase or water (1 μL in 25 μL) for 2 h was measured for at least three dilutions in the linear calibration range. All data was acquired using the default Luminscence application in a SpectraMax iD3 (Molecular Devices). See supplementary material for more details.

### Protein and nucleic acid quantification

To estimate the protein content of our Xenopus Extract, we employed the BioRad DC Protein Assay with the standard 15 minutes incubation time and without using the surfactant solution. The samples absorbance was measured in disposable 1 cm optical path cuvettes at 750 nm using an Amersham Bioscience Ultrospec 2100 Pro Spectrophotometer. Upon calibration using BSA (MilliporeSigma), the typical concentration of proteins in our preparation was found to vary between 40 ∼ 45 g/L among different batches, comparable to reported estimates of the cytoplasmic concentration of non-yolk proteins in Xenopus eggs (*36*). Nucleic acids were quantified from UV absorbance as 260 nm after removing the protein contribution, using an extinction coefficient at 260 nm equal to 40 μg/mL.

## Acknowledgments

the authors would like to thank Kehui Xiang for his support in developing the Xenopus extract preparation, Elinor Ng Eaton for her support in using the SpectraMax iD3 and Kelsey Farenhem for assistance in the analysis of DMS reactivity data.

## Funding

This work is supported by grants from the NIH (R35GM151111), Chan Zuckerberg Initiative (DAF2022-250422), Richard and Susan Smith Family Foundation, and the David and Lucile Packard Foundation.

## Competing interests

Authors declare that they have no competing interests.

## Notes

### Competing Interest Statement

The authors have declared no competing interest.

