## supplementary materials for "A passive protein environment offsets RNA folding energetics in cells"

### 1. List of sequences and constructs

| Name | Sequence (5′ → 3′) |
|---|---|
| Surface Tether | /Biotin/ TTTT GAGG CTTA ACGG ACCG |
| Hairpin Acceptor | CAGC CAA /Cy5/ G CCAG GTAT CGGT CCGT TAAG CCTC |
| HP5 Stem Loop | ATAC CTGG CTTG GCTG TT rGrCrGrGrC TTTT TTTT TTTT TTTT TTTT TTTT TTTT TT rGrCrCrGrC TT /Cy3/ |
| HP5d Stem Loop | ATAC CTGG CTTG GCTG TT GCGGC TTTT TTTT TTTT TTTT TTTT TTTT TTTT TT GCCGC TT /Cy3/ |
| HP5r Stem Loop | rArUrArC rCrUrGrG rCrUrUrG rGrCrUrG rUrU rGrCrGrGrC rUrUrUrU rUrUrUrU rUrUrUrU rUrUrUrU rUrUrUrU rUrUrUrU rUrUrUrU rUrU rGrCrCrGrC rUrU TT /Cy3/ |
| HP8 Stem Loop | ATAC CTGG CTTG GCTG TT rArGrCrArGrUrGrC TTTT TTTT TTTT TTTTT TTTT TTT rGrCrArCrUrGrCrU TT /Cy3/ |
| HP10a Stem Loop | ATAC CTGG CTTG GCTG TT rCrUrGrCrArGrUrGrUrC TTTT TTTT TTTTT TTTT TTT rGrArCrArCrUrGrCrArG TT /Cy3/ |
| HP10b Stem Loop | ATAC CTGG CTTG GCTG TT rCrGrArGrCrArGrUrGrC TTTT TTTT TTTTT TTTT TTT rGrCrArCrUrGrCrUrCrG TT /Cy3/ |
| HP10c Stem Loop | ATAC CTGG CTTG GCTG TT rCrGrCrCrArGrCrUrGrC TTTT TTTT TTTTT TTTT TTT rGrCrArGrCrUrGrGrCrG TT /Cy3/ |
| HP6 Stem Loop | ATAC CTGG CTTG GCTG TT rCrGrArGrCrC TTTTTTTTTT rGrGrCrUrCrG TT /Cy3/ |
| HP9 Stem Loop | ATAC CTGG CTTG GCTG TT rArCrArUrGrArCrUrArT TTTTTTTTT rUrArGrUrCrArUrGrU TT /Cy3/ |
| HP5 L5 Stem Loop | ATAC CTGG CTTG GCTG TT rGrCrGrGrC TTTTT rGrCrCrGrC TT /Cy3/ |
| HP4 Stem Loop | ATAC CTGG CTTG GCTG TT rGrCrArC TTTTTTTTTTTTTT rGrUrGrC TT /Cy3/ |
| H2A L4 | ATAC CTGG CTTG GCTG TT rGrGrCrUrCrU TTTT rArGrArGrCrC TT /Cy3/ |
| H2A L10 | ATAC CTGG CTTG GCTG TT rGrGrCrUrCrU TTTTTTTTTT rArGrArGrCrC TT /Cy3/ |
| H2A L20 | ATAC CTGG CTTG GCTG TT rGrGrCrUrCrU TTTTTTTTTT TTTTTTTTTT rArGrArGrCrC TT /Cy3/ |
| RNA Duplex Acceptor | /Cy3/ TT **rCrArGrCrCrUrA** rUrGrArCrG TT CGGT CCGT TAAG CCTC |
| RNA Duplex Donor | **rUrArGrGrCrUrG** /Cy5/ |
| DNA Duplex Donor | /Cy5/ **GGCTTGG** |
| RNA-5 nt | rCrCrUrUrG |
| RNA-8 nt | rUrUrGrGrArUrUrA |
| RNA-10 nt | rCrCrUrUrGrGrArUrUrA |
| RNA-16 nt | rUrUrUrCrUrUrCrCrUrCrUrUrUrCrCrU |
| RNA-30 nt | rUrUrCrUrCrUrArGrUrCrUrGrGrCrGrGrArGrGrUrGrGrCrUrGrArUrUrUrA |
| RNA-61 nt | rGrUrUrCrArGrArGrUrUrCrUrArCrArGrUrCrCrGrArGrGrArCrCrUrGrArGrArArArGrGrGrUrCrGrGrArGrArUrCrGrGrArArGrArGrCrArCrArCrGrUrCrU |
| RNA-103 nt | rGrGrArUrCrGrArArArGrArUrUrUrUrUrUrCrUrUrCrCrGrArGrGrGrArGrGrArGrArCrGrUrArGrGrCrArCrCrUrArCrArGrUrGrCrGrArArArGrCrArCrGrUrGrCrArCrCrArCrCrGrUrUrCrUrCrArGrCrArCrGrUrArCrCrCrGrArArGrArUrCrGrGrArArGrArGrCrCrC |

**Table S1.** List of all sequences used to perform experiments in this work. Matching color/font styles highlight base-pairing regions across sequences. The letter “r” is used to signify the insertion of a ribonucleotide in the sequence.

| Construct | Components |
| --- | --- |
| HP5 | Surface Tether + Hairpin Acceptor + HP5 Stem Loop |
| HP5d | Surface Tether + Hairpin Acceptor + HP5 Stem Loop |
| HP5r | Surface Tether + Hairpin Acceptor + HP5r Stem Loop |
| HP8 | Surface Tether + Hairpin Acceptor + HP8 Stem Loop |
| HP10a | Surface Tether + Hairpin Acceptor + HP10a Stem Loop |
| HP10b | Surface Tether + Hairpin Acceptor + HP10b Stem Loop |
| HP10c | Surface Tether + Hairpin Acceptor + HP10c Stem Loop |
| HP6 Stem Loop | Surface Tether + Hairpin Acceptor + HP6 Stem Loop |
| HP9 Stem Loop | Surface Tether + Hairpin Acceptor + HP9 Stem Loop |
| HP5 L5 Stem Loop | Surface Tether + Hairpin Acceptor + HP5 L5 Stem Loop |
| HP4 Stem Loop | Surface Tether + Hairpin Acceptor + HP4 Stem Loop |
| H2A L4 | Surface Tether + Hairpin Acceptor + H2A L4 Stem Loop |
| H2A L10 | Surface Tether + Hairpin Acceptor + H2A L10 Stem Loop |
| H2A L20 | Surface Tether + Hairpin Acceptor + H2A L20 Stem Loop |
| RNA duplex | Surface Tether + RNA Duplex Acceptor + RNA Duplex Donor |
| DNA duplex | Surface Tether + Hairpin Acceptor + DNA Duplex Donor |

**Table S2.** List of constructs and relative components used in this work.

#### 2. Assessment of hairpin behavior with Xenopus extract

In this work we have used the folded fraction (θ) of a surface-tethered hairpin as a reporter for the effect of the Xenopus extract on nucleic acid thermodynamics. Our design of choice involved an RNA stem embedded in a DNA construct, as commonly done in the field of sm-tFRET.
To make sure that the behavior observed was not due to the specific design choices, we validated our assay for full-DNA (HP5d) and full-RNA (HP5r) constructs, finding the same qualitative behavior (Fig. S1).

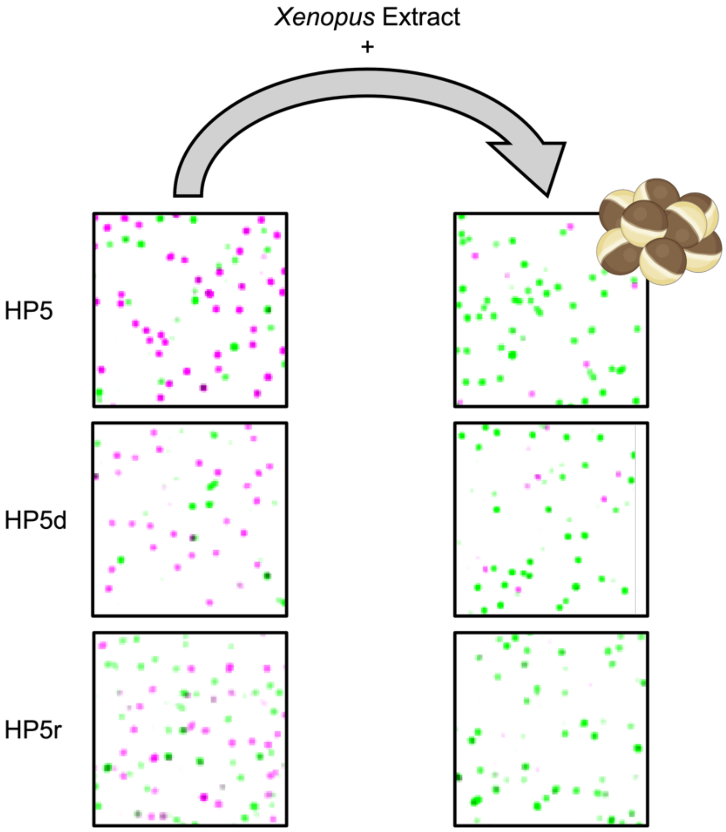

**Fig. S1.** Effect of Xenopus extract on hairpin with different design choice. HP5 has an RNA stem embedded in a DNA construct, while HP5d is a full DNA construct and HP5r is a full RNA construct. All of them are unfolded by Xenopus extract.

#### 3. Tethering the construct does not alter its folding

To test whether tethering our construct was affecting its folding equilibrium, we performed a series of melting measurements with the exact same construct free in solution.
Folding free energy at room temperature was extrapolated from the melting curves, finding good agreement with our sm-tFRET results (see Fig S2).

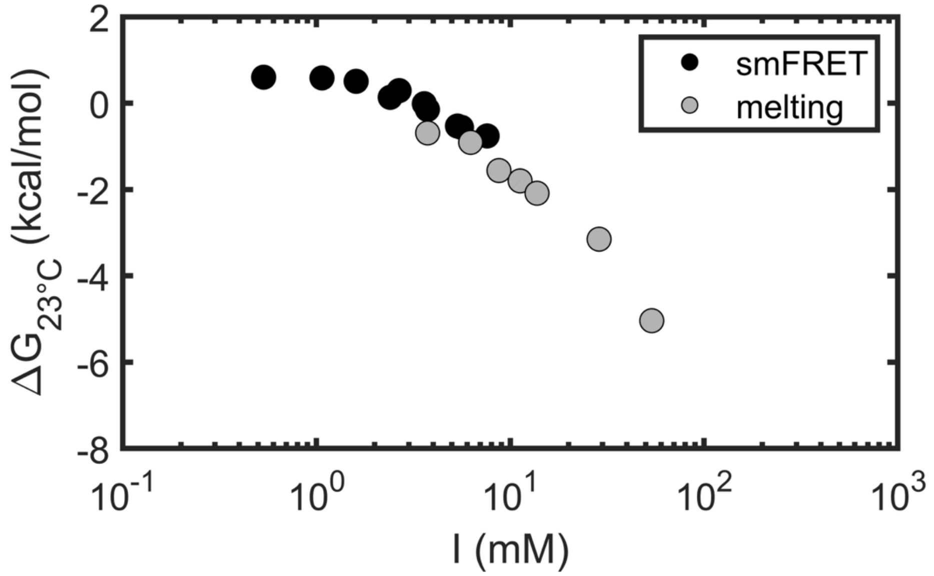

**Fig. S2.** Tethering (sm-tFRET) doesn't affect the folding equilibrium when compared to the behavior extrapolated from melting measured in solution.

#### 4. Xenopus extract causes loss of dynamic behavior

Adding Xenopus Extract to HP5 causes hairpins to be promptly unfolded. At the same time, the dynamic opening and closing of the hairpins is completely loss, making the system apparently static.

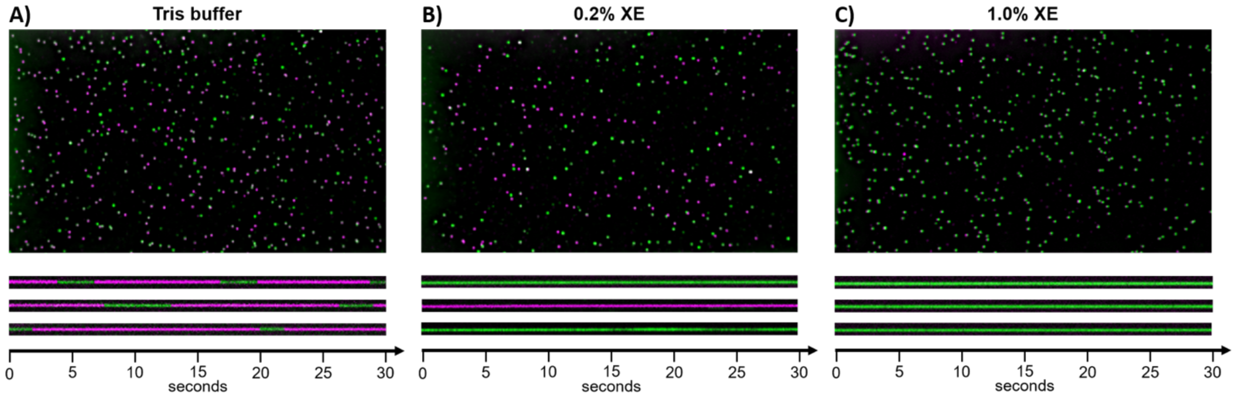

**Fig. S3.** Representative snapshots of HP5 (top) and spectrographs (bottom) in pure buffer (A) and increasing concentration of XE, 0.2% (B) and 1.0% (C). The spectrographs show the color of three selected particles as a function of time. In buffer, the particles swap between folded and unfolded state. When XE is added, no dynamic behavior can be measured over the typical sm-tFRET experimental timescales.

#### 5. ATP quantification and Apyrase treatment

To evaluate whether the effect of Xenopus Extract was active (ATP-dependent) or passive (ATP-independent) we have treated the extract with Apyrase. To avoid interference with the provided buffer, we have directly added the enzyme to our extract and then verified that it was performing as expected and consuming all available ATP.
To quantify the amount of ATP in the extract, we used the ATP-Assay Kit-Luminesce from Dojindo (product A550). The assay reports on Luciferase activities and relies on the sample's ATP to fuel the reaction and convert D-Luciferin to Oxyluciferin, emitting visible light as a byproduct. We generated a calibration curve to map Luciferase Intensity ($x$) to ATP concentration by fitting the following polynomial function (Fig. S4A):

$$[ATP](x) = Ax^3 + Bx^2 + Cx$$

We then measured the light emitted by the extract for at least three dilutions that yielded values in the linear range of the calibration curve with or without Apyrase treatment. We found that the concentration of ATP in the raw extract was equal to (2.8 ± 0.4) mM and upon Apyrase treatment it was instead equal to (2.6 ± 3.3) μM, virtually indistinguishable from background noise (Fig. S4B). This simple analysis confirms the effectiveness of the enzymatic treatment and corroborates the unfolding activity of the Xenopus Extract as being passive (ATP-independent).

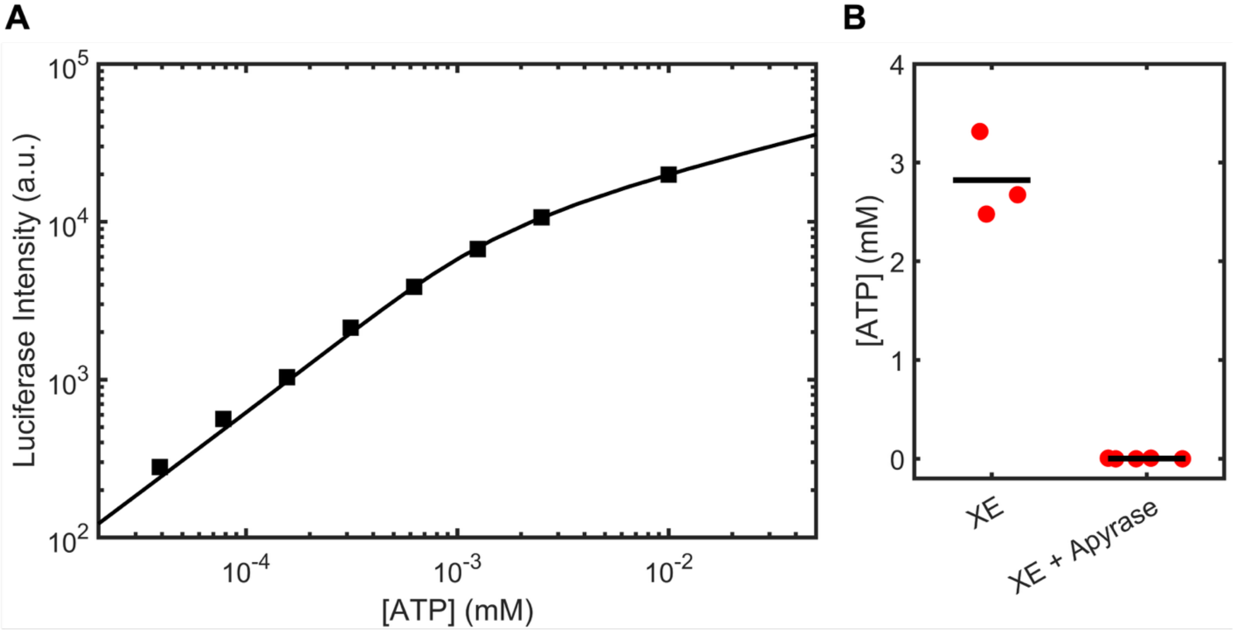

**Fig. S4.** Quantification of ATP in the Xenopus Extract before and after Apyrase treatment. A) Calibration curve for the ATP-Assay Kit-Luminesce from Dojindo. B) Quantified ATP concentration in XE before and after enzymatic treatment, confirming the efficacy of Apyrase in degrading virtually all available ATP.

### 6. Calibration of the apparent salt concentration in *Xenopus* extract

To fit our model, we must input the folding energy of our hairpins ($\Delta G_F$), and we opted to use calculated values from the well-established NN model for nucleic acids (1). One caveat with this approach is that the NN values are given at 1 M NaCl, which is not appropriate for our system. When our system is at zero XE, its ionic strength is extremely small and contributed mostly by the Tris buffer. As the concentration of XE in our system increases, the ionic strength increases by an unknown amount. This means that $\Delta G_F$ itself is a function of the concentration of XE.

*Xenopus* extract is a complex mixture containing ions, small molecules and a large variety of macromolecules. To disentangle the effect of the ionic composition and the larger molecular components on the behavior of nucleic acids, we fractionated the extract using an Amicon filter 3K and studied the effect of the flowthrough ("light fraction").
We found that the light fraction stabilizes nucleic acid interactions, as expected when increasing the amount of salt in solution. To quantify this effect, we measured the hybridization of one RNA duplex construct (see Table S1 and S2 for exact sequences) as we titrated against salt or the XE light fraction.
The extent of hybridization was measured with a SpectraMax iD3 monitoring the emission of the Cy3 donor fluorophore. Binding curves at different RNA concentrations were subsequently analyzed to extract dissociation constants (K).
We observed that the logarithm of the dissociation constants determined this way scaled linearly with the logarithm of the concentration of either salt or XE light fraction, so that:

$$\begin{cases} \ln[K] = \ln[NaCl] \cdot m_{NaCl} + q_{NaCl} \\ \ln[K] = \ln[XE_{LF}] \cdot m_{XE_{LF}} + q_{XE_{LF}} \end{cases}$$

By solving this set of equations, we could conveniently derive a mapping function that links the Light Fraction concentration to NaCl concentration (Fig. S5):

$$\ln[NaCl] = \frac{\ln[XE_{LF}] \cdot m_{XE_{LF}} + q_{XE_{LF}} - q_{NaCl}}{m_{NaCl}}$$

$$\ln[NaCl] = \frac{\ln[XE_{LF}] \cdot m_{XE_{LF}}}{m_{NaCl}} + \frac{q_{XE_{LF}} - q_{NaCl}}{m_{NaCl}}$$

$$\ln[NaCl] = \ln\left([XE_{LF}]^{m_{XE_{LF}}/m_{NaCl}}\right) + \frac{q_{XE_{LF}} - q_{NaCl}}{m_{NaCl}}$$

$$[NaCl] = [XE_{LF}]^{m_{XE_{LF}}/m_{NaCl}} \cdot e^{\frac{q_{XE_{LF}} - q_{NaCl}}{m_{NaCl}}}$$

$$[NaCl] = 0.22 \cdot [XE_{LF}]^{1.43}$$

Using this calibration, it becomes possible to map an effective NaCl concentration to any given XE volume fraction. This way, we found that the pure light fraction of XE behaves as a ~160 mM NaCl solution, and our calibration curve was used to correct the values of $\Delta G_F$ at any XE concentration (2). Having an estimate for $\Delta G_F$, we were left with only one unknown variable, $\Delta G_{XE}$ (kcal/mol), that could be fit to our data.

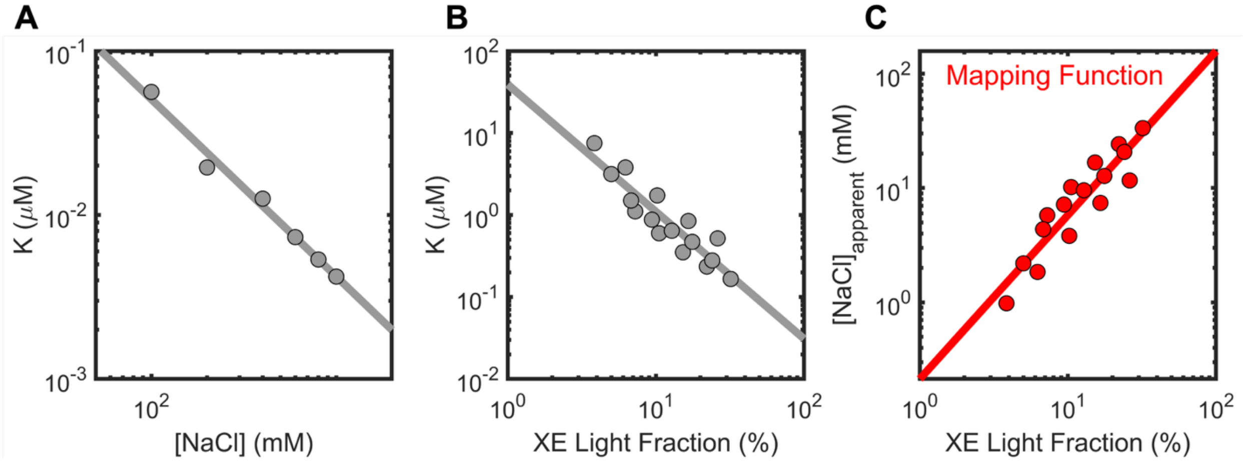

**Fig. S5.** Calibration of Xenopus Extract apparent salt concentration. A) Dissociation constant for one RNA duplex in buffer (10mM Tris pH 8.0) and added NaCl. B) Dissociation constant of the same RNA duplex in Xenopus Extract Light Fraction. C) Mapping function converting XE Light Fraction to NaCl.

### 7. Derivation of model

The protein shows a length-dependence in its binding. If an oligonucleotide has *N* independent binding sites (so that the binding footprint is equal to *RNA length / N*), we can take advantage of the binomial theorem to write the RNA folded fraction as follows:

$$\theta = \frac{e^{-\Delta G_f * \beta}}{e^{-\Delta G_f * \beta} + \left(1 + e^{-\Delta G_p * \beta}\right)^N}$$

With $\beta = 1/RT$, with RT being 0.58 kcal/mol.
We can divide both numerator and denominator by the binomial term:

$$\theta = \frac{e^{-\Delta G_f * \beta} / \left(1 + e^{-\Delta G_p * \beta}\right)^N}{e^{-\Delta G_f * \beta} / \left(1 + e^{-\Delta G_p * \beta}\right)^N + 1}$$

Since $\left(1 + e^{-\Delta G_p * \beta}\right)^N$ can be written as $e^{N * ln\left(1 + e^{-\Delta G_p * \beta}\right)}$, we have that:

$$\theta = \frac{e^{\left[-\Delta G_f - N * RT * ln\left(1 + e^{-\Delta G_p * \beta}\right)\right]\beta}}{e^{\left[-\Delta G_f - N * RT * ln\left(1 + e^{-\Delta G_p * \beta}\right)\right]\beta} + 1}$$

We can conveniently rewrite $-RT * ln\left(1 + e^{-\Delta G_p * \beta}\right)$ as an apparent protein binding energy $\Delta G_p'$ per binding site. In the strong binding regime ($e^{-\Delta G_p * \beta} \gg 1$), which holds for the binding energies relevant here, $\Delta G_p' \longrightarrow \Delta G_p$.

$$\theta = \frac{e^{\left(-\Delta G_f + N * \Delta G_p'\right)\beta}}{e^{\left(-\Delta G_f + N * \Delta G_p'\right)\beta} + 1}$$

Making RNA length explicit we have that $N * \Delta G_p' = RNA\ length * \frac{\Delta G_p'}{\text{footprint}}$.
Because the footprint and the per-site energy enter only through their ratio, the two parameters cannot be determined independently from our data, so we absorb them into a single per-nucleotide destabilization $\Delta G_p^{nt} \equiv \frac{\Delta G_p'}{\text{footprint}}$, giving an exponent $RNA\ length * \Delta G_p^{nt}$:

$$\theta = \frac{e^{\left(-\Delta G_f + RNA\ length * \Delta G_p^{nt}\right)\beta}}{e^{\left(-\Delta G_f + RNA\ length * \Delta G_p^{nt}\right)\beta} + 1}$$

Meaning that the three-state competition behaves as two-state folding equilibrium, with the hairpin stability lowered by a fixed amount per nucleotide, and $RNA\ length * \Delta G_p^{nt}$ thus behaving as an effective destabilization term.
